# Optimizing the live attenuated influenza A vaccine backbone for high-risk patient groups

**DOI:** 10.1101/2021.10.06.462766

**Authors:** João P.P. Bonifacio, Nathalia Williams, Laure Garnier, Stephanie Hugues, Mirco Schmolke, Beryl Mazel-Sanchez

## Abstract

The live attenuated influenza vaccine (LAIV) is approved for intranasal spray application in 2-49 year-old patients with safety concerns limiting its use in younger children and immunocompromised patients, mainly from the higher incidence of adverse events and the possibility of uncontrolled replication and reversion to a pathogenic strain, respectively. Further attenuation of the LAIV could generally improve its safety profile, which might come at the cost of reduced immunogenicity. To solve this dilemma, we took advantage of a recently defined mechanism of ER stress induction by modifying IAV non-structural protein 1 (NS1). The modified LAIV (AAmut/PR8) showed stronger ER stress activation *in vitro* and replicated to lower titers *in vivo* compared to its parental strain, without affecting protection against homo-subtypic or hetero-subtypic IAV strains. AAmut/PR8 could pose as a suitable strategy to attend the gap to the current LAIV recommendation guidelines in susceptible target populations.

## Introduction

Influenza A viruses (IAV) are the etiological agents causing one of the most significant respiratory tract infections. Hospitalizations and deaths occur mainly among high-risk groups, which include pregnant women, children under 5-year-old, elderly and immunocompromised patients. In the context of the recent pandemic, influenza related hospitalizations and deaths have been lower than previous years^1–3^, most likely due to preventive measures widely adopted such as hand washing, sanitizing, keeping distances and providing better air ventilation. However, seasonal IAV infections are still an important threat to healthcare systems as a recent metadata analysis showed 7% of total acute lower respiratory infection cases in children under 5 years were attributed to IAV^4^. Additionally, an out-of-season resurgence in flu infections is expected, as the unexposed population grows wider during the pandemic^5^.

One of the most effective ways to prevent influenza diseases is vaccination. Three types of vaccines are available for IAV: recombinant influenza vaccines or split-virus inactivated influenza vaccine (IIV), both injected intra-muscularly and a live-attenuated influenza vaccine (LAIV), in the form of a nasal spray. The LAIV uses the genetic backbone of a temperature sensitive (*ts*), cold-adapted (*ca*), attenuated (*att*) influenza strain derived from A/Ann Arbor/6/1960 restricting vaccine virus replication to the upper respiratory tract at 33 °C, thus preventing disease onset. Additionally, LAIV mimicks a natural infection, which brings the advantage of triggering mucosal immune responses^6^. Hence, the general vaccine effectiveness (VE) of the LAIV is as good if not better in young children than the VE of the inactivated vaccine^7,8^. Adversely, its safety profile excludes vaccination of high-risk patients. Current guidelines advise against the use in immunocompromised patients, pregnant women, adults over 50 years old and infants under 2 years old. In fact, higher rates of fever, rhinorrhea and wheezing after LAIV administration were shown in the latter population^9,10^. Therefore, different strategies have been explored to improve the safety of the LAIV with a large majority of them targeting the IAV non-structural protein 1 (NS1)^11–15^.

NS1 is a virulence factor that modulates the cellular anti-viral response^16,17^ and interferes with host mRNA maturation by interacting with the 30 kDa subunit of the cleavage and polyadenylation specificity factor (CPSF30)^18,19^. CPSF30 is a key factor in the poly-adenylation complex and in its absence, pre-mRNAs are not polyadenylated and thus rapidly degraded, leading to a reduced host cell protein production^20^. The interaction between NS1 and CPSF30 is well described in the literature with a crystal structure showing the amino acids responsible for this interaction. While positions 103 and 106 are important to stabilize the interaction^21,22^, positions 108, 125 and 189 were shown to antagonize the innate immune response by reducing antiviral genes expression^23^.

We recently demonstrated that CPSF30 binding by NS1 is essential for blocking the unfolded protein response (UPR)^24^. Importantly, the UPR contributes to the adjuvant effect of the licensed adjuvant AS03^25^, which is a component of inactivated influenza vaccine Prepandrix. In our study, we proposed a targeted approach to mutate NS1 from the LAIV backbone and render it unable to bind CPSF30. We hypothesized that by modulating LAIV’s capacity of inducing a host response, we could improve its safety profile with more efficient viral clearance and limited viral replication in the host, whilst maintaining its ability to induce a protective immune response from the activation of ER stress and UPR pathway. If successful, this approach could widen the patient spectrum in which a live attenuated vaccine could be applied.

## Results

### Abrogation of CPSF30 binding does not diminish production of LAIV in ovo or in MDCK cells

Deletion of the expression of the non-structural protein 1 (NS1) from LAIV has been shown to increase its attenuated phenotype^13,15^. We attempted a targeted approach to attenuate the LAIV by abolishing the ability of NS1 to interact with CPSF30. We used targeted mutagenesis to substitute F103S, M106I, K124E, D144R and D189G from the LAIV *ca* backbone and rescued the mutant virus (AAmut/PR8) by reverse genetics. The NS1 mutant virus and its parental counterpart (AAwt/PR8) are composed of the six gene segments of the cold-adapted influenza A/Ann Arbor/6/1960 plus the segments of hemagglutinin and neuraminidase of influenza A/Puerto Rico/8/1934 (PR8) and their only difference is in its NS1 gene sequence. Both viruses were grown in embryonated chicken eggs at 33 °C and sequenced to confirm the presence of the mutations. To confirm that the five aa mutations introduced abrogate the NS1-CPSF30 binding, we overexpressed a flag-tagged CPSF30 protein in HEK 293T cells and superinfected those cells with AAmut/PR8 or AAwt/PR8. Following co-immunoprecipitation, AAwt/PR8 NS1 was able to bind CPSF30 while AAmut/PR8 NS1 showed only residual binding (Figure 1a).

**Figure 1:**
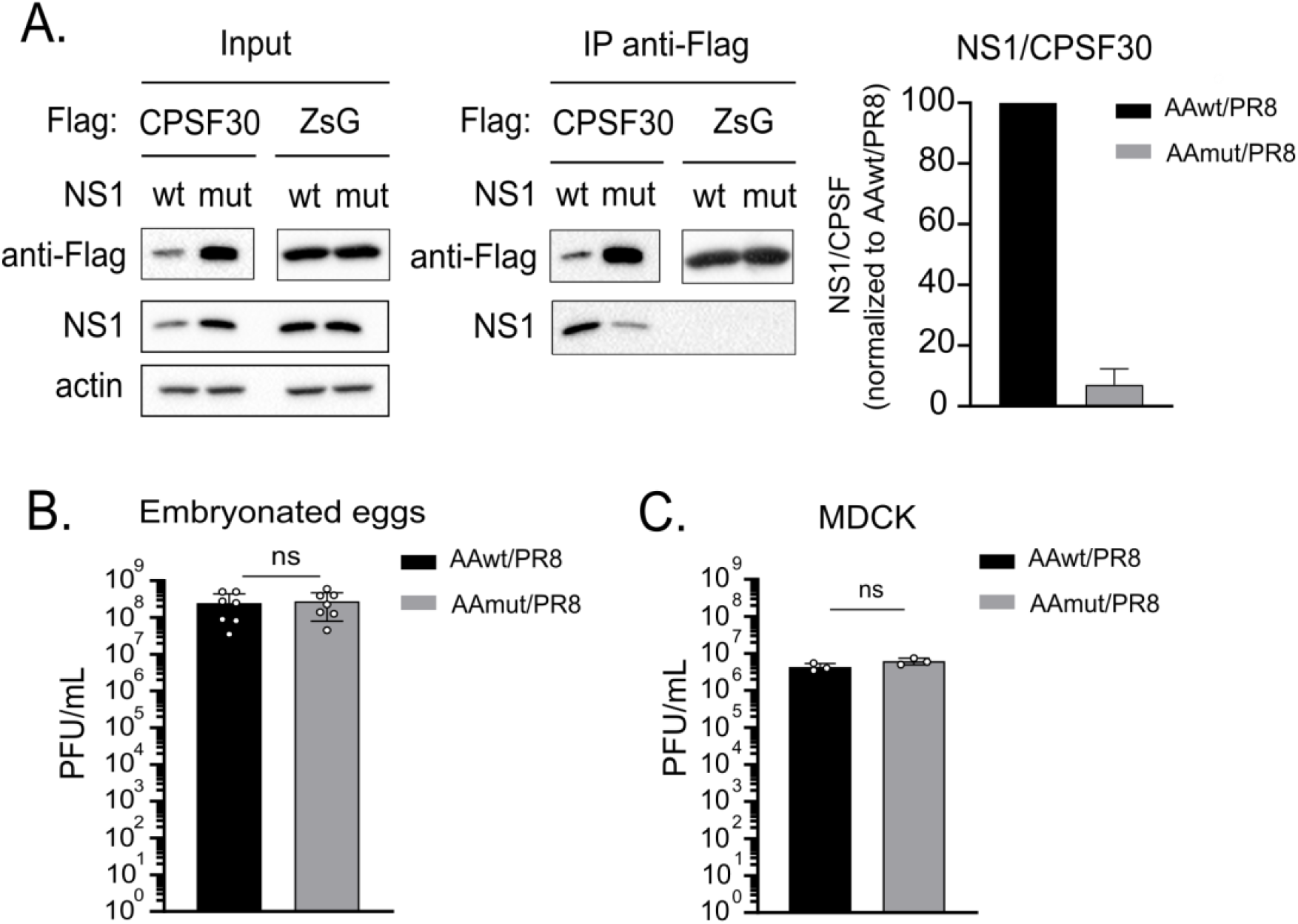
AAmut/PR8 NS1 binds to CPSF30 without compromising vaccine virus growth in eggs or MDCK cells. (a) 293T cells were transfected with pCAGGS.Flag-CPSF30 or pCAGGS.Flag-ZsGreen for 24 h prior infection with either AAwt/PR8 or AAmut/PR8 at a MOI of 5. Cells were incubated for 16 h at 33 °C and then lysed. Anti-Flag M2 affinity beads were used to immunoprecipitate (IP) the flag-tagged protein and its interactors. Precipitates were separated by SDS-PAGE and detected by WB using anti-flag antibody and anti-NS1 antibody and the amount of NS1 protein bound to Flag-CPSF30 was quantified using densiometric analysis (n=2) (b and c) Ten-days old embryonated chicken eggs (b) were infected with 100 PFU and MDCK B cells (c) were infected at a MOI of 0.1. After 46 h at 33 °C allantoic fluid and supernatant were collected. Viral titers were determined by plaque assay in MDCK cells at 33 °C. Graphs represent the mean ± SD of n=7 repeats in eggs or n=3 repeats in MDCK cells, respectively, and the statistical significances were determined using unpaired t-test. *ns* = non-significant

Enhanced attenuation of LAIV could result in deleterious replication defects, with unwanted consequences for large scale vaccine production. We thus compared the replication of AAmut/PR8 and AAwt/PR8 in two relevant production models, embryonated chicken eggs and MDCK cells. Titers of AAwt/PR8 and AAmut/PR8 vaccine stock were comparable in both production systems (Figure 1b, c).

These results indicate that the targeted mutagenesis of LAIV significantly impaired its CPSF30 binding without affecting the potential for vaccine production.

### AAmut/PR8 is attenuated without compromising protection in an adult mouse model

In order to demonstrate that our strategy results in an attenuated phenotype *in vivo*, we first compared the replication of AAmut/PR8 and AAwt/PR8 in the upper and lower respiratory tract of adult mice. At 4 days post-vaccination we consistently detected AAwt/PR8 virus in snouts and lungs, while AAmut/PR8 was absent in all snout samples and present in only two lung samples at lower viral loads compared to AAwt/PR8 (Figure 2a, b). Since AAmut/PR8 displayed a clear replication defect, we asked, if the protection elicited by AAmut/PR8 was compromised compared to AAwt/PR8. 8-weeks old mice were vaccinated intranasally with 10^2^-10^4^ pfu. The highest dose corresponds approximately to the recommended weight adjusted dose of quadrivalent LAIV in human adults^26^. We challenged the animals at 21 days post-vaccination with 10 LD_50_ of influenza A/Netherlands/602/2009 (H1N1) strain (homosubtypic/heterologous challenge), simulating a severely drifted virus. We monitored body weight (Figure 2c) and survival (Figure 2d) for 14 days. All animals were protected after either vaccine (Figure 2d) when using 10^4^ pfu. While we found a trend towards reduced protection with lower doses of AAmut/PR8 (Figure S1a, b), there was overall no statistically significant difference observed for the protective dose 50 (PD_50_) calculated for the dose response to both vaccines (Figure 2e). We also showed that neither vaccine caused any alterations in body weight of adult mice (Figure 2f), reflecting the already attenuated phenotype of this virus. In conclusion, compared to AAwt/PR8, our vaccine candidate AAmut/PR8 had lower replication in adult’s mice respiratory tract, while conferring full protection against a homosubtypic/heterologous challenge.

**Figure 2:**
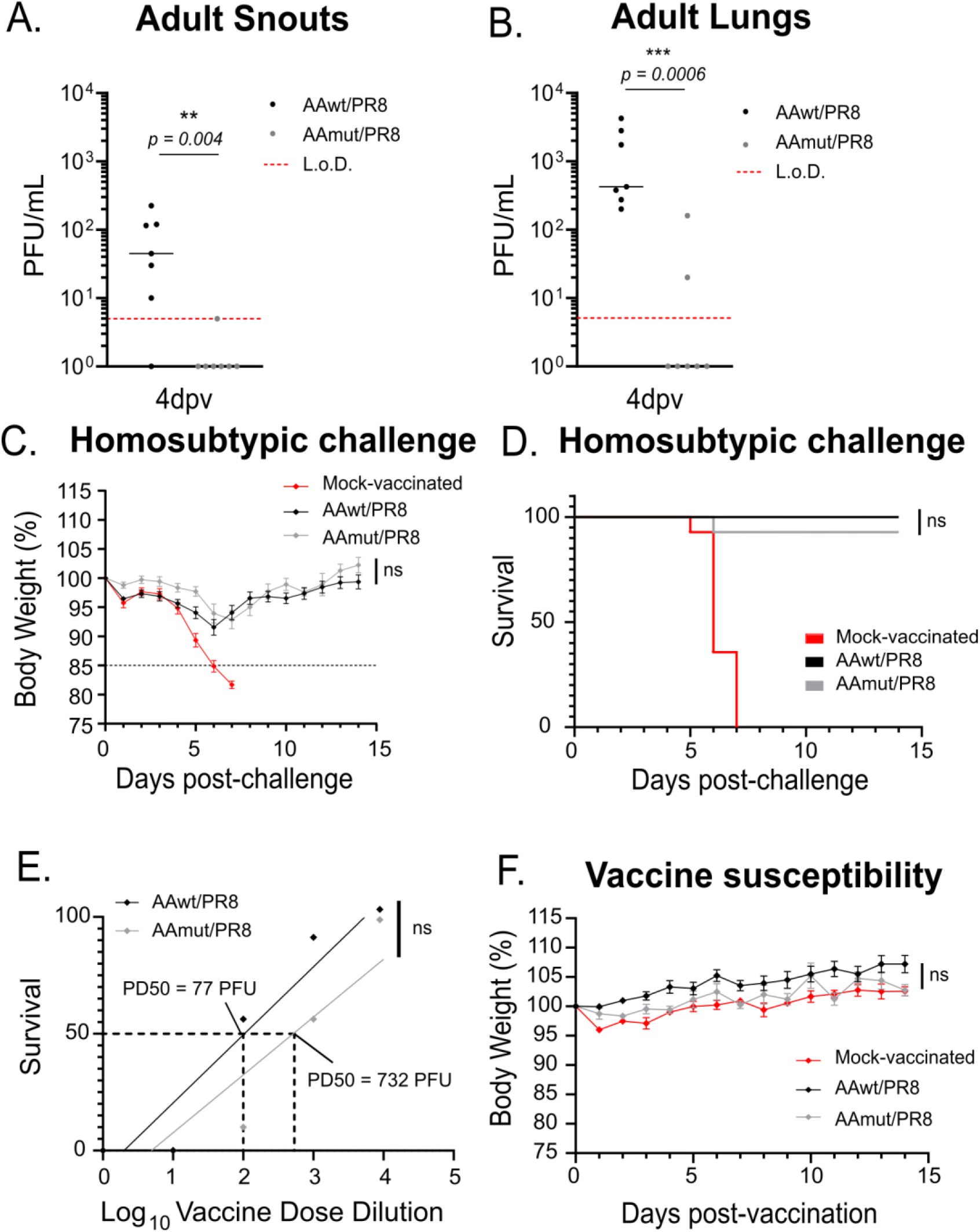
AAmut/PR8 is attenuated but protects adult mice against a homosubtypic/heterologous challenge. (a and b) Female 8-weeks-old mice were vaccinated intranasally under anaesthesia with 10^5^ PFU AAwt/PR8 or AAmut/PR8 in 25 μL PBS. At 4 days post-vaccination snouts (a) and lungs (b) were collected and vaccine viral titters were determined by plaque assay (n=7 per group). (c - e) Female 8-weeks-old mice vaccinated with 10^4^ PFU of AAwt/PR8 or AAmut/PR8 were challenged at day 21 post-vaccination with 20 PFU (10×LD_50_) of mouse adapted A/Netherlands/602/2009 (H1N1) in 20 μL PBS under light anaesthesia (n=14 per group). Body weight (c) and survival (d) were monitored for 14 days post-challenge. PD_50_ was calculated according to Reed & Muench. Linear regression was used to determine statistical significance between the two PD_50_ (e). (f) Body weight was followed for 14 days after vaccination with 10^4^ PFU of AAwt/PR8 or AAmut/PR8. The statistical significances between AAwt/PR8 group and AAmut/PR8 group were determined using Mann-Whitney test in panels a and b; two-way ANOVA with the Geisser-Greenhouse correction and post-hoc Dunn’s multiple comparisons test in panel c and f and Mantel-Cox test in panel d. *\*p < 0*.*05*, \*\**p < 0*.*01*, \*\*\**p < 0*.*001, ns* = non-significant, L.o.D. = limit of detection (5 PFU/mL). Graphs are representative of 3 independent experiments and indicate median for panels a and b or mean ± SEM for panels c and f. Symbols represent data from individual mice for panels a and b. Black dotted line represents 15% body weight loss cut-off.

### AAmut/PR8 is attenuated without compromising protection in neonatal mice

Younger children were reported to show adverse effects, such as wheezing after vaccination^9^. We postulated that by showing lower replication, our vaccine candidate has a higher attenuation degree, which could ameliorate the safety concerns regarding its use in infants younger than 2 years. Thus, we characterized AAwt/PR8 and AAmut/PR8 replication in a more appropriate model using 7-days old mice. We vaccinated neonatal mice via the intranasal route and determined vaccine viral loads after 2, 4 and 6 days. In accordance with the results obtained in adult mice, at 2 days post-vaccination we observed lower titers of AAmut/PR8 compared to AAwt/PR8 in snouts (Figure 3a). While at day 4 both vaccine viral loads were similar and reduced compared to day 2, we observed that on day 6 post-vaccination, AAwt/PR8 virus titers were dispersed with some animals presenting high titers while others had non-detectable virus. This was not the case for the AAmut/PR8 group, which at day 6 post-vaccination had either low viral titers or non-detectable virus. In the lungs, we did not consistently detect virus after infection with either vaccine, which might be a consequence of vaccine administration in absence of anesthesia. We conclude from these data that AAmut/PR8 is cleared faster from the airways of neonatal mice compared to AAwt/PR8. It is well known that the immune system of young children is in an immature state with compromised immune responses to vaccines^27,28^. Thus, we asked if our attenuation phenotype would have an impact on the protection elicited by this vaccine in our neonatal model. To answer this question, we evaluated both short-term protection after a heterosubtypic challenge and long-term protection after a homosubtypic/heterologous challenge. Neonatal mice were vaccinated with increasing doses of each vaccine and challenged with either influenza A/Vietnam/2004 after 21 days or A/Netherlands/09 after 49 days. In the heterosubtypic challenge, significant protection was observed in all doses for both vaccines in a similar fashion (Figure 3b,c and Figure S2a-b). Interestingly, in the long-term homosubtypic/heterologous challenge, while the middle dose of AAmut/PR8 was not protective compared to AAwt/PR8 (Figure S2c-d), both vaccines were similarly protective at the higher dose of 10^4^ PFU (Figure 3d,e). Additionally, there was no overall statistically significant difference observed for the protective dose 50 (PD_50_) of each vaccine (Figure 3f). We also showed that neither vaccine caused any reduction in body weight gain of neonatal mice (Figure 3g), once again reflecting the already attenuated phenotype of this virus. In conclusion, compared to AAwt/PR8, our vaccine candidate AAmut/PR8 is mildly attenuated in neonatal mice, while still able to protect against a long-term homosubtypic challenge and a heterosubtypic challenge.

**Figure 3:**
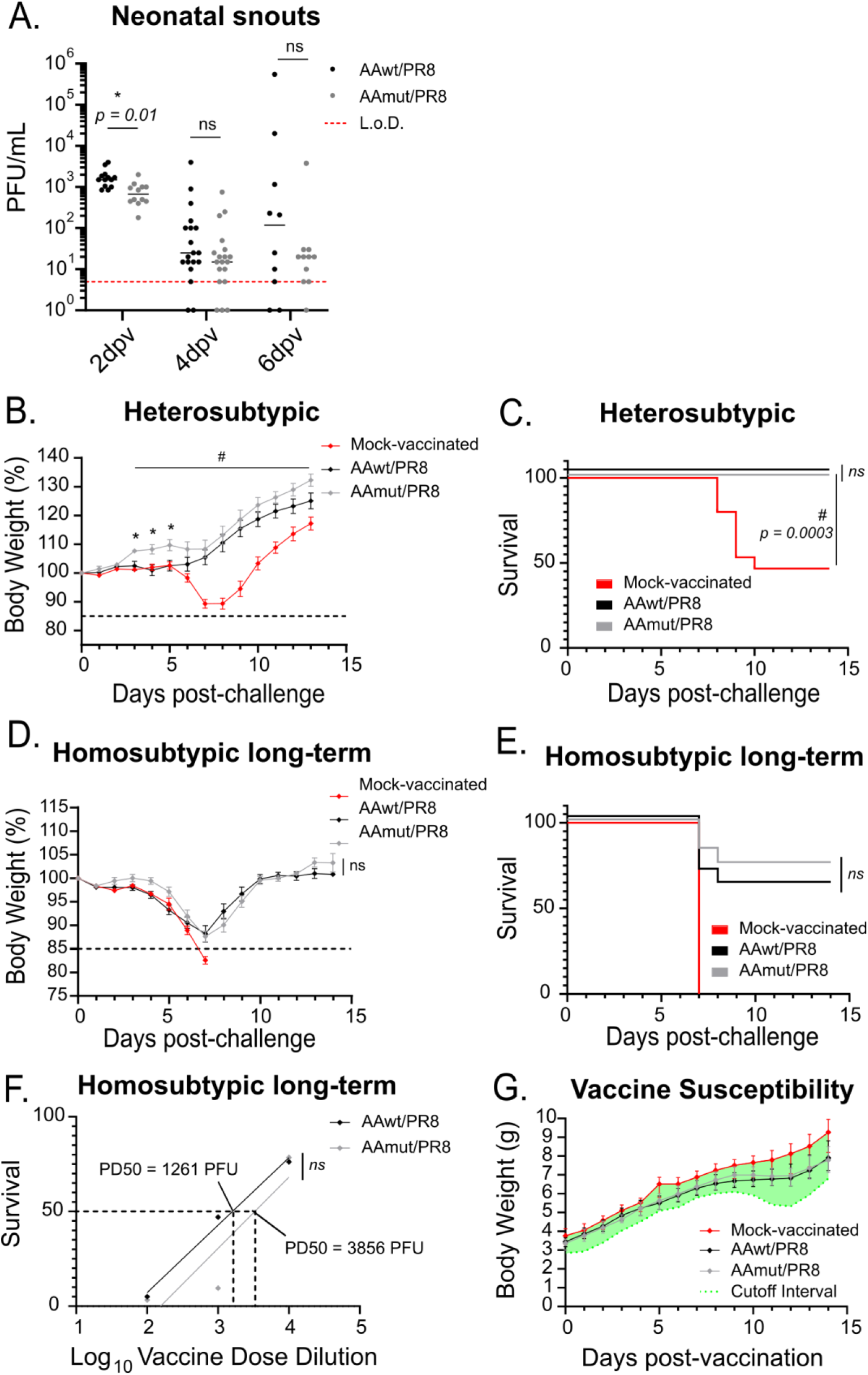
AAmut/PR8 is attenuated and protects neonatal mice against long-term homosubtypic/heterologous challenge and heterosubtypic challenge. (a) Seven-days-old mice were vaccinated intranasally with 10^5^ PFU of AAwt/PR8 or AAmut/PR8 in 5 μL PBS. At 2, 4 and 6 days post-vaccination vaccine viral titters in the snouts were determined by plaque assay (n=10-19 per group). (b – c) Vaccinated 7-days-old mice with 10^4^ PFU of AAwt/PR8 or AAmut/PR8 were challenged at day 21 post-vaccination with 10^3^ PFU (20×LD_50_) of A/Vietnam/1203/2004 (H5N1) in 20 μL PBS under anaesthesia (n=11 per group). Body weight (b) and survival (c) were monitored for 14 days post-challenge. (d – f) Vaccinated 7-days-old mice with 10^4^ PFU of AAwt/PR8 or AAmut//PR8 were challenged at day 49 post-vaccination with 20 PFU (10×LD_50_) of A/Netherlands/602/2009 (H1N1) in 20 μL PBS under anaesthesia (n=11-17 per group). Body weight (d) and survival (e) were monitored for 14 days post-challenge. PD_50_ was calculated according to Reed & Muench. Linear regression was used to determine statistical significance between the two PD_50_ (f). (g) Body weight was followed for 14 days after vaccination with 10^4^ PFU of AAwt/PR8 or AAmut/PR8 and the cut-off was defined as twice the difference between the average weight of the non-vaccinated group and the lightest mouse of that group, calculated daily. The statistical significances between AAwt/PR8 group versus AAmut/PR8 group were determined using two-way ANOVA with the Geisser-Greenhouse correction and post-hoc Dunn’s multiple comparisons test in panels a, b and d and Mantel-Cox test in panel c and e. *\*p < 0*.*05*, \*\**p < 0*.*01*, \*\*\**p < 0*.*001*, ^***#***^*p < 0*.*05* between mock-vaccinated and AAmut/PR8 groups, *ns* = non-significant, L.o.D. = limit of detection (5 PFU/mL). Graphs are representative of 2-3 independent experiments and indicate median for panel a, mean ± SEM for panels b, d and g. Symbols represent data from individual mice for panel a.

### AAwt/PR8 and AAmut/PR8 induces overall comparable innate immune responses in neonatal mice snouts

Blocking of CPSF30 by NS1 is a cornerstone in the host protein synthesis shutoff initiated by IAV infection. This broadly active strategy limits the innate antiviral response^29^. To further explore the safety profile of AAmut/PR8, we looked into the innate response induced by both vaccines in neonatal mice snouts at 2 days post-vaccination, when vaccine loads were highest. Both vaccines induced antiviral cytokines *Cxcl10* and *Ccl5* expression at similar levels indicating similar innate responses. When we looked into pro-inflammatory cytokines, no significant expression of *Tnfa* and *Il6* was observed compared to mock-vaccinated animals (figure 4). Overall, our data do not support a differential induction of innate immune responses as a reason for the enhanced attenuation while maintaining immunogenicity.

**Figure 4:**
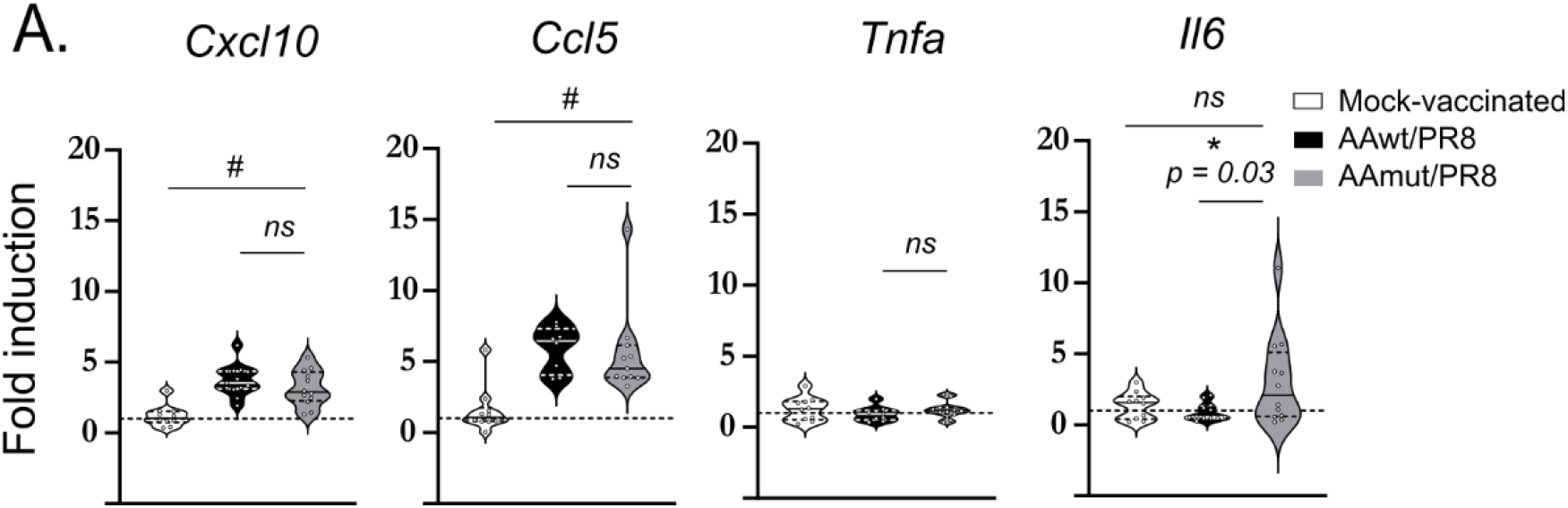
AAwt/PR8 and AAmut/PR8 induces overall comparable innate immune responses in neonatal mice snouts. Seven-days-old mice (n=11-12) were vaccinated intranasally with 10^5^ PFU of AAwt/PR8 or AAmut/PR8 in 5 μL PBS. At 2 days post-vaccination snouts were harvested and RT-qPCR was performed from isolated RNA. The statistical significances between groups were determined using one-way ANOVA and post-hoc Tukey’s multiple comparisons test. *\*p < 0*.*05*, \*\**p < 0*.*01*, \*\*\**p < 0*.*001*, ^***#***^*p < 0*.*05 ns* = non-significant. Graphs are representative of 2 independent experiments and median is represented.

Remarkably, *Il6* expression was suppressed in AAwt/PR8 vaccinated animals compared to AAmut/PR8 (figure 4). The same NS1 mutations introduced in AAmut/PR8 NS1 were described to modulate ER stress activation and UPR through the XBP1 pathway^24^ which can lead to IL-6 expression ^30,31^. Therefore, to explore this pathway as a mechanism of attenuation we evaluated the expression of two target genes regulated by the UPR. Neither AAmut/PR8 nor AAwt/PR8 vaccinated neonatal and adult mice induced significant expression of *Dnajb9* (ERdj4) or *Ddit3* (CHOP) compared to mock-vaccinated animals (Figure S3). UPR is notoriously difficult to measure *in vivo* due to its transient activation and a poor signal-to-noise ratio from a low percentage of infected cells in the upper respiratory tract (refs). Thus, we decided to characterize AAmut/PR8 ability to modulate ER stress activation using relevant *in vitro* cell models.

### AAmut/PR8 induces ER stress in human and mouse cells

NS1 binding to CPSF30 was previously shown to modulate the induction of the UPR^24^. Therefore, we asked if the mutation in AAmut/PR8 NS1 allowed the establishment of the UPR upon infection. We infected A549 cells and evaluated XBP1 splicing and sXBP1 protein expression levels by western blot. Our results show an increase in the spliced form of XBP1 protein levels in AAmut/PR8-infected A549 as compared to AAwt/PR8 (Figure 5a). In addition, the spliced form of XBP1 mRNA amount observed by semi-quantitative RT-PCR matched with the increased amount of spliced XBP1 protein (Figure 5b). We also analysed by RT-qPCR the induction of genes downstream of the transcription factor sXBP1, namely *DDIT3* and *DNAJB9*. Compared to AAwt/PR8, the expression of these two genes was increased in cells infected with AAmut/PR8 (Figure 5c, upper panel), in line with previous published results. This increase in *Dnajb9* and *Ddit3* was also observed when we used a murine lung epithelial cell line (Figure 5c, lower panel) indicating that this phenomenon is species independent and reflects a general phenotype during viral replication, when the ability of NS1 to bind CPSF30 is abrogated. Also, these observations were not explained by a difference in replication of AAmut/PR8 since viral replication was comparable in both cell models (Figure 5d).

**Figure 5:**
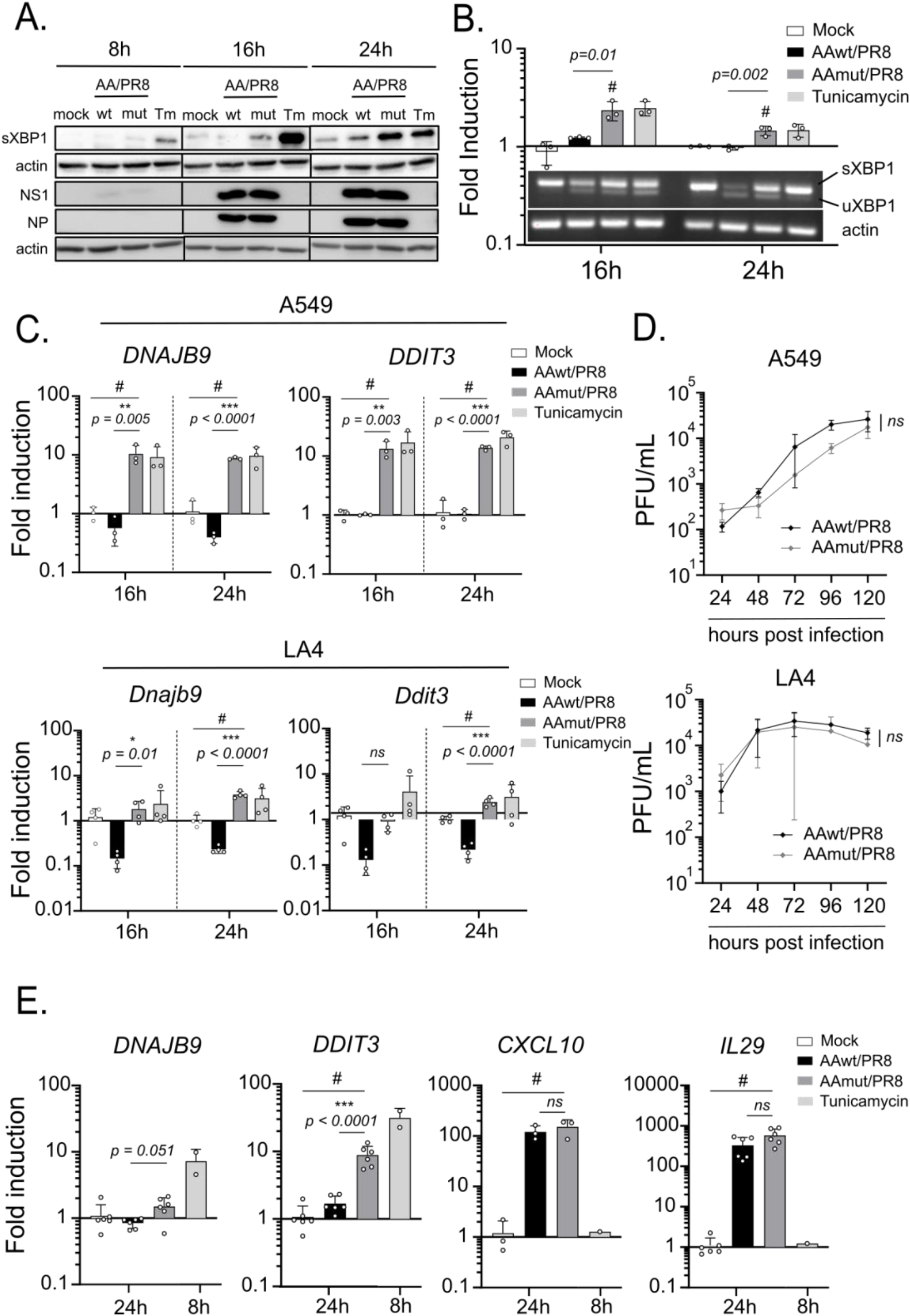
AAmut/PR8 NS1 allows unfolded protein response activation in murine and human cell models. (a - b) A549 cells were infected at a MOI of 5 with AAwt/PR8 or AAmut/PR8 or treated with tunicamycin at 5 ug/ml (n=3). Cells were collected at 8, 16 and 24 h post infection and lysates were analysed by western blot for sXBP1, actin, NP and NS1 protein levels (a). At 16 h and 24 h post infection RT-PCR was performed from isolated RNA for the spliced form of XBP1 mRNA (b). Graph represents fold induction of sXNP mRNA. (c) A549 cells (upper panel) or LA4 cells (lower panel) were infected at a MOI of 5 with AAwt/PR8 or AAmut/PR8 or treated with tunicamycin at 5 ug/ml (n=3). Cells were collected at 16 and 24 h post infection RT-qPCR for UPR-induced genes DNAJB9 and DDIT3. (d) A549 (upper panel) or LA4 (lower panel) were infected at a MOI of 0.01 with either AAwt/PR8 or AAmut/PR8. Supernatants were collected at indicated time post infection and viral titer determined by plaque assay. (e) Primary human nasal epithelial cells (Mucilair) were infected at a MOI of 5 with AAwt/PR8 or AAmut/PR8 (n=6) or treated with tunicamycin (n=2) at 5 ug/ml for 8 h. Cells were lysed at 24 h post infection and RT-qPCR for each respective gene was performed. The statistical significances between AAwt/PR8 group versus AAmut/PR8 group were determined using one-way ANOVA with post-hoc Tukey’s multiple comparison test for panels b-d and g and two-way ANOVA with the Geisser-Greenhouse correction and post-hoc Dunn’s multiple comparisons test for panels e and f. *p < 0.05, **p < 0.01, ***p < 0.001, ^**#**^p < 0.05 between mock and AAmut/PR8 groups; ns = non-significant. Graphs are representative of 2-3 independent experiments and indicate mean ± SD.

Lastly, we took advantage of a more relevant model using stratified primary human nasal epithelial cells grown in air-liquid interface culture. These cells would be the first encountering the LAIV virus in case of vaccination of human patients. In this model, we also observed induction of both UPR-induced genes and no differences in the induction of antiviral response genes (Figure 5e).

Overall, our results show that AAwt/PR8 and AAmut/PR8 mount a comparable antiviral response in relevant cell models and *in vivo*, which do not explain the reduced *in vivo* replication and comparable immunogenicity. In contrast, AAmut/PR8 allows the establishment of a robust UPR compared to AAwt/PR8 *in vitro*, which might contribute to the enhanced attenuated profile of AAmut/PR8 observed *in vivo*.

## Discussion

LAIV is currently only approved for use in target population ranging from healthy 2 years-old to 49 years-old. Thus key-populations, such as pregnant women, elderly subjects, children under 2 years-old and immunocompromised patients do not benefit from the advantages of this vaccine option, such as a wider immune response and avoidance of traumatic injections specially in paediatrics. The rationale behind this limitation is the fear of reversion to a virulent strain that could cause disease based on experience with other replicating vaccines such as measles-mumps-rubella (MMR), yellow fever and oral polio, which have shown reported cases of uncontrolled replication and were able to cause disease in immunocompromised patients^32–35^. Not surprisingly, several attempts have tried to improve LAIV safety, including approaches that either delete the NS1 gene of LAIV or mutate it, rendering a more attenuated virus ^11–15^. In fact, two clinical trials tested NS1-deleted-NS1 LAIVs containing HA and NA from influenza virus H1N1, H3N2, and B strains in adult volonteers. As for the parental LAIV a single intranasal dose was well tolerated, safe, not shedded from the patient and induced significant levels of strain-specific and cross-neutralizing antibodies in vaccinated humans^36,37^. However, it remains currently unclear, if a similar safety profile could be achieved in more susceptible patient groups.

Another major concern in administering LAIV in children under 2 years-old is the higher incidence of side effects observed such as wheezing, cough and runny nose. Although there is a lack of evidence pinpointing to the mechanism responsible for this, an association between vaccine virus shedding and runny nose after vaccination in this target population has been described^38^. We showed here lower viral replication and earlier clearance of a CPSF30 binding mutant LAIV in neonatal mice, which hints to a generally safer profile in children under 2 years old.

The innate immune response to a replicating virus contributes to the onset of the symptoms associated with LAIV administration in this population. Several cytokines and chemokines have been associated to lung damage during acute influenza infection. Indeed, CXCL-10, RANTES, IL-6 and TNF-a are all involved in the host response to IAV and can lead to a detrimental response^39,40^. A potential danger of eliminating the antagonism of CPSF30 in LAIV could have been an exaggerated inflammatory response. However, in our in vivo models, AAmut/PR8 induced similar levels of pro-inflammatory cytokines as AAwt/PR8 suggesting that the attenuated phenotype observed does not exacerbate the host immune response to the vaccine.

Curiously, we observed a slightly lower expression of Il-6 in AAwt/PR8 vaccinated neonatal mice (Figure 4). The pathophysiological role of this cytokine is still unclear, with reports showing a correlation between its expression and clinical score symptoms in human patients^39,41^ while others report a protective role for this cytokine during influenza infection in mice^42–44^. Moreover, IL-6 has been described as a target gene of the UPR transcription factor sXBP1 during ER stress activation^30,45,46^. *In vitro*, AAmut/PR8 more potently induces ER stress activation and UPR in accordance with our previous work^24^. Unfortunately, we could not demonstrate this pathway’s activation *in vivo* most likely due to the sensitivity of the techniques used. The lack of appropriate models such as IRE1α or XBP1 knockout mice also renders it difficult to assess ER stress activation impacts *in vivo*.

Notwithstanding, together with the fact that AAmut/PR8 is attenuated and still able to protect both adult and neonatal mice against a homosubtypic and heterosubtypic challenge, we propose that ER stress activation during AAmut/PR8 administration compensates for its lower replication and maintains this vaccine immunogenicity and efficacy. This hypothesis is further corroborated by the fact that ER stress has been directly implicated in the establishment of an immune response to influenza vaccines as a mechanism of action of the adjuvant AS03^25^.

The work shown here characterizes for the first time the LAIV virus in a neonatal mice model and presents an alternative vaccine candidate to LAIV with an even more pronounced attenuated phenotype while still able to maintain its protection. We confirm that this vaccine is still scalable and able to be produced at similar titers as the current vaccine virus used in Fluenz® Tetra to attend a population-wise demand and presents a safer option to the current LAIV with the possibility of being administered to target populations where uncontrolled replication of the virus is a serious concern.

