## Supplemental figures for "Optimizing the live attenuated influenza A vaccine backbone for high-risk patient groups"

FIG S1

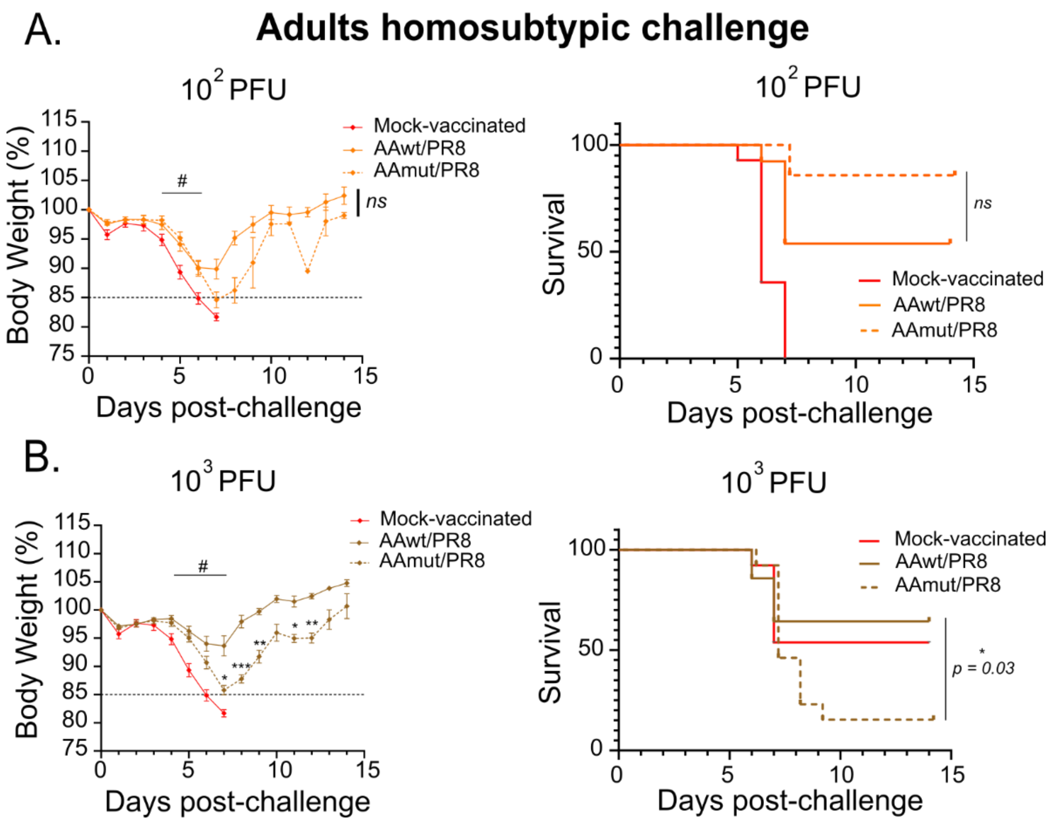


**Figure S1: AAwt/PR8 and AAmut/PR8 body weight loss and survival in adult mice after challenge.** Female 8-weeks-oldmice vaccinated with (a) $10^2$ PFU or (b) $10^3$ PFU of either AAwt/PR8 or AAmut/PR8 were challenged at day 21 post-vaccination with 20 PFU ($10 \times LD_{50}$) of A/Netherlands/602/2009 (H1N1) in 20 μL PBS under anaesthesia (n=14 per group).

Body weight (left panels) and survival (right panels) were monitored for 14 days post-challenge. The statistical significances between AAwt/PR8 group versus AAmut/PR8 group were determined using two-way ANOVA with the Geisser-Greenhouse correction and post-hoc Dunn's multiple comparisons test for panels a and b. **p < 0.05, **p < 0.01, ***p < 0.001,* $^{\#}p < 0.05$ between mock and AAmut/PR8 groups; *ns* = non-significant. Graphs are representative of 2 independent experiments and indicate mean ± SEM.

FIG S2

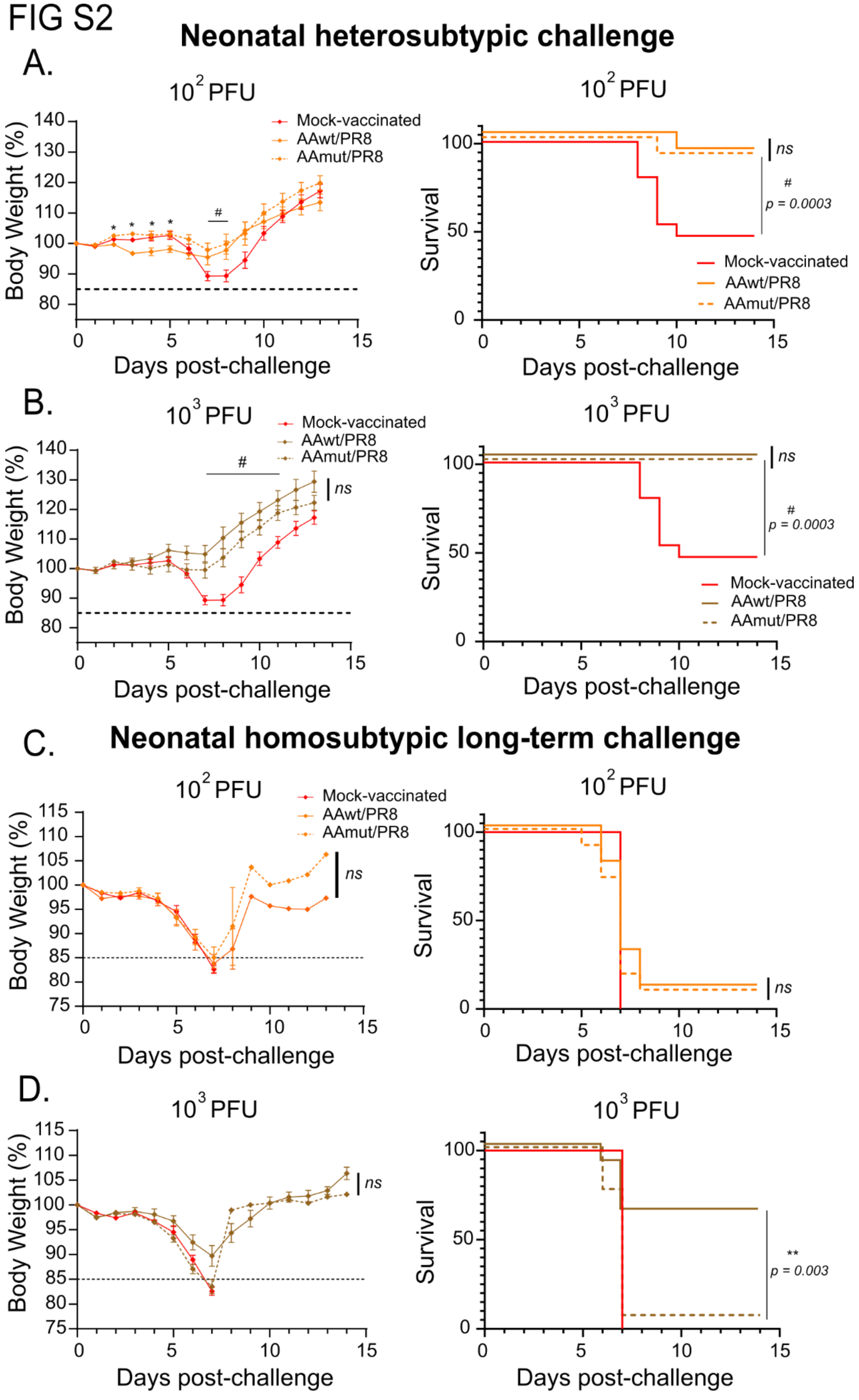


**Figure S2: AAwt/PR8 and AAmut/PR8 body weight loss and survival after challenge in neonatal mice.**

(a-b) Seven-days-old mice were vaccinated intranasally with $10^2$ PFU (a) or $10^3$ PFU (b) of AAwt/PR8 or AAmut/PR8 in 5 $\mu$L PBS. At day 21 post-vaccination, mice were challenged with $10^3$ PFU (20×$LD_{50}$) of A/Vietnam/1203/2004 (H5N1) in 20 $\mu$L PBS under anaesthesia (n=11 per group). Body weight (left panels) and survival (right panels) were monitored for 14 days post-challenge.

(c-d) Seven-days-old mice were vaccinated intranasally with $10^2$ PFU (c) or $10^3$ PFU (d) of AAwt/PR8 or AAmut/PR8 in 5 $\mu$L PBS. At day 49 post-vaccination, mice were challenged with 20 PFU (10×$LD_{50}$) of A/Netherlands/602/2009 (H1N1) in 20 $\mu$L PBS under anaesthesia (n=11-17 per group). Body weight (left panels) and survival (right panels) were monitored for 14 days post-challenge.

The statistical significances between AAwt/PR8 group versus AAmut/PR8 group were determined using two-way ANOVA with the Geisser-Greenhouse correction and post-hoc Dunn's multiple comparisons test for panels a,b and d,e. *$p < 0.05$, **$p < 0.01$, ***$p < 0.001$, $^{\#}p < 0.05$ between mock-vaccinated and AAmut/PR8 groups *ns* = non-significant. Graphs are representative of 2 independent experiments and indicate mean ± SEM.

FIG S3

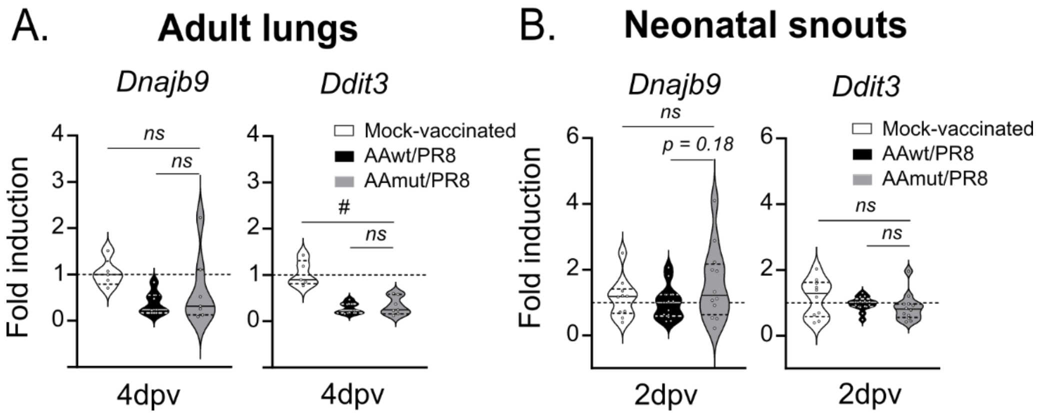


**Figure S3: UPR induced genes after AAwt/PR8 and AAmut/PR8 vaccination in adult lungs and neonatal snouts.**

(a) Female 8-weeks-old mice were vaccinated intranasally under anesthesia with $10^5$ PFU of AAwt/PR8 or AAmut/PR8 in 25 μL PBS. At 4 days post-vaccination lungs were harvested and RT-qPCR performed from isolated RNA for UPR-induced genes *Dnajb9* and *Ddit3*.

(b) Seven-days-old mice(n=11-12) were vaccinated intranasally with $10^5$ PFU of AAwt/PR8 or AAmut/PR8 in 5 μL PBS. At 2 days post-vaccination snouts were harvested and RT-qPCR performed in isolated RNA for UPR-induced genes *Dnajb9* and *Ddit3*.

The statistical significances between AAwt/PR8 group versus AAmut/PR8 group were determined using one-way ANOVA and post-hoc Tukey's multiple comparisons test. $*p < 0.05$, $**p < 0.01$, $***p < 0.001$, $^{\#}p < 0.05$ between mock-vaccinated and AAmut/PR8 groups *ns* = non-significant. Graphs are representative of 2 independent experiments and indicate media.
